# Improved management facilitates return of an iconic fish species

**DOI:** 10.1101/197780

**Authors:** Brian R. MacKenzie, Kim Aarestrup, Kim Birnie-Gauvin, Massimiliano Cardinale, Mads Christoffersen, Henrik S. Lund, Iñigo Onandia, Gemma Quilez-Badia, Mark R. Payne, Andreas Sundelöf, Claus Sørensen, Michele Casini

**Affiliations:** National Institute for Aquatic Resources (DTU Aqua), Technical University of Denmark, Kemitorvet, Building 150, DK 2800 Kongens Lyngby, Denmark; National Institute for Aquatic Resources (DTU Aqua), Technical University of Denmark, Silkeborg, Denmark; Swedish University of Agricultural Sciences, Department of Aquatic Resources, Institute of Marine Research, Turistgatan 5, Lysekil 45330, Sweden; Danish Fishermen Producer Organisation, Nordensvej 3, Taulov, DK 7000 Fredericia, Denmark; AZTI Tecnalia, Marine Research Division, Herrera Kaia Portualdeaz/g, 20110 Pasaia, BasqueCountry, Spain; Catalan Association for Responsible Fishing, Barcelona, Catalonia, Spain; Marinexperten, H.C.Ørstedsvej 9, DK 9900 Frederikshavn, Denmark

**Keywords:** bluefin tuna, range, recovery, prey, management

## Abstract

Species declines and losses of biota are often associated with shifting baselines in perceived historical abundances, and/or neglect or abandonment of recovery actions aimed at ecological restoration. Such declines are frequently accompanied by contractions in the geographical distribution of the species, with associated negative ecological impacts and diminishing socio-economic benefits. Here we show using citizen science and other data that after 50-60 years of near total absence, the iconic top predator and highly migratory species bluefin tuna, *Thunnus thynnus*, returned by the hundreds if not thousands in waters near Denmark, Norway and Sweden during August-October 2015-2017. The re-utilisation of this former habitat is part of a geographically more widespread expansion of the summer foraging area to the northern part of the northeast Atlantic Ocean, encompassing waters from east Greenland to west Sweden. The remarkable return to the Skagerrak, Kattegat and North Sea has been facilitated by improved fishery management for bluefin tuna and its prey. Bluefin tuna biomass in the northeast Atlantic and Mediterranean has been increasing since a recovery plan was implemented in the late 2000s, and biomasses of two key prey species (herring, *Clupea harengus*; mackerel, *Scomber scombrus*) recovered during the late 1980s-1990s. The reappearance of bluefin tuna in the Skagerrak-Kattegat and other waters of northern Europe, despite a recent history of mismanagement and illegal fishing in the northeast Atlantic and Mediterranean which led to a critical population decline, offers hope that other marine ecological recoveries are possible under improved management of fisheries and ecosystems.

**One Sentence Summary:** Improved management promotes the return of an ocean icon to northern Europe.

**Significance Statement:** Commercial fisheries are often perceived being in a state of decline and collapse, putting food and economic security at risk. Such declines are frequently accompanied by contractions in stock distribution, negative ecological impacts and diminishing socio-economic benefits. Here we present an example based on one of the world’s most valuable and iconic fish species, bluefin tuna, which demonstrates that effective management of both bluefin tuna and its prey has been a key factor leading to a remarkable reoccupation of formerly lost habitat. This reappearance, following decades of absence, occurred despite the bluefin tuna stock having had a recent, long history of unsustainable and illegal exploitation. Marine ecological recovery actions can be successful, even in situations which may initially appear intractable.

## Introduction

Recovering the biomass and spatial range of depleted fish stocks is challenging for fisheries managers and conservation ecologists (1–5). Once lost, former biomasses and ranges often disappear from human memory, thereby reducing motivation for and impairing recovery efforts (6–8). Biomass and range recovery often take longer than anticipated, even when fishing is reduced below sustainable levels, due to several factors such as bycatch fishing mortality (i. e. captures as bycatch in other fisheries), and changes in stock productivity (1–4, 9, 10). Species that are highly migratory and whose migrations take them into the high seas and multiple fishing jurisdictions, such as tunas and billfishes, are potentially even more vulnerable to depletion and prolonged recovery times than stocks under single or few fishery jurisdictions (2, 11).

Here we report a recovery of the former range of distribution by a highly migratory, highly valuable, iconic top predator fish species (bluefin tuna, *Thunnus thynnus*). This species was the target of unsustainable exploitation for many years in the 1990s-2000s, and its biomass declined to record low levels in the late 2000s (12). As one of the world’s most valuable fish species, it has also been the subject of much scientific, public and conservation scrutiny (13).

Bluefin tuna spawn in sub-tropical regions (e.g., Mediterranean Sea) and then migrate north long distances as adults to summer and autumn foraging areas (14, 15). In the northeast Atlantic, historical adult foraging areas are located in the Bay of Biscay, on the northwest European continental shelf off Ireland and the UK, and in the Norwegian Sea-North Sea-Skagerrak-Kattegat-Øresund (the latter are hereafter referred to as northern European waters; Supplementary Figure S1 showing ICES areas and sea names (14, 16)). Bluefin tuna occupy these waters in summer-early autumn before migrating southwards to overwintering areas. This long-distance migratory behaviour is a part of the species’ life-history, having evolved through generations (15).

However, the migrations to northern European waters stopped almost completely in the early-mid-1960s and bluefin tuna have been extremely rare in the area since the 1970s (16, 17); the species has not supported targeted commercial or recreational fisheries in the area since then (12) (Supplementary Figure S2). The reasons for both the disappearance and the long period before reappearance are unclear, but likely due to a combination of overexploitation of juveniles and adults of both the tuna and their prey, and changing oceanographic conditions (16, 18–20).

Similar large changes in range of fish stocks have been reported in the past for some other fish species as populations decline and expand (3, 4, 21, 22). Theoretical and conceptual models attribute such changes to density – dependent effects on habitat utilisation, biological rates and properties such as growth rates, habitat availability and cannibalism, as well as social interactions and collective memory within populations (19, 21, 23).

In this report, we describe the re-appearance of bluefin tuna in northern European waters using citizen science data (i. e., observations from non-scientists pursuing activities on or near the sea) and our own observations, and we discuss possible reasons why the species has returned to this region. Given the high level of illegal and unsustainable exploitation of this species in other parts of its range in the recent past (mid-late 1990s until ca. 2008-2010; (12)), the recovery of the habitat and range of this species is extraordinary and could become a classic example of how recoveries can occur under a suitable combination of fishery management regulations and ecosystem conditions.

## Results

### Bluefin tuna observations in 2015-2017

We found and received many reports of bluefin tuna in the region. The observations are summarized in Figure 1 and Supplementary Table S2. The reports included observations of single individual tunas and of schools of various sizes from a few specimens (2–10) to hundreds. The species is relatively easy to identify and distinguish in this area, mainly because of their surface jumping behaviour, body shape, color and size. The sizes of tuna observed were usually large (ca. 2 – 3 m, corresponding approximately to ages 8+, based on length-age relationships for this population) and the jumping or surface-breaking behaviour is characteristic of this species. Most observations we present are based on individuals, which are partially or entirely out of the water due to jumping and surface swimming, which facilitated reliable identification; the observations are supported in many cases by photographs or videos available on public social media and angling or news media websites in Scandinavia. Some of these photographs are shown in Figure 2 and as Supplementary Figure S3, and some links to online videos of bluefin tuna jumping are listed in Supplementary Table S3. Other observations occurred from a bluefin tuna tagging study conducted in the area in September 2017 (24).

**Figure 1.**
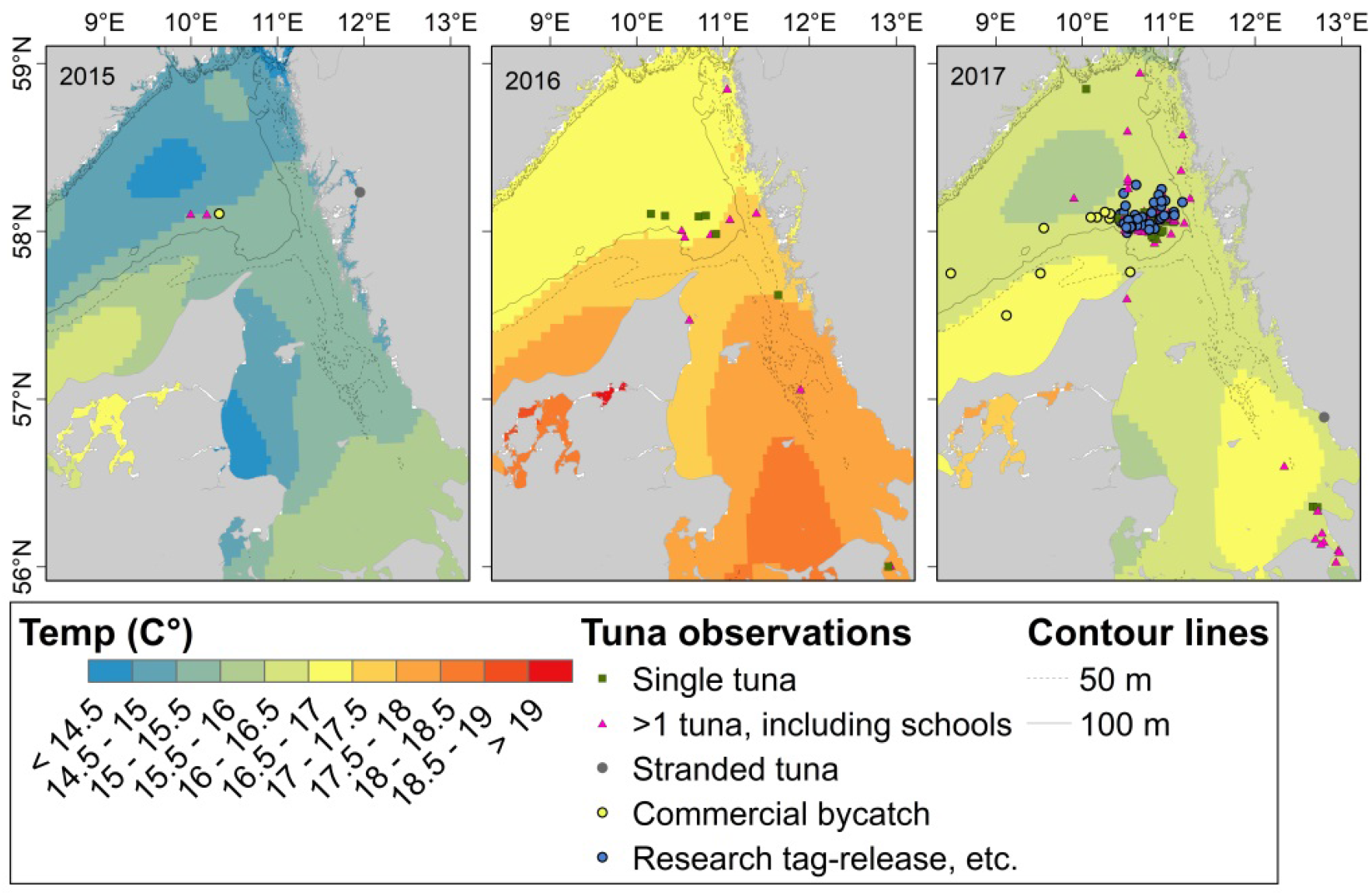
Locations where individual or schools of bluefin tuna were observed in Skagerrak-Kattegat-Øresund in 2015-2017. Colour contours: averaged sea surface temperature as derived from satellite imagery (64) during Sept. 7-21 in each of the three years. Black contour lines: bottom topography. During 2017, bluefin tuna were also captured for tagging purposes and in commercial fisheries as bycatches. These catches are indicated with different symbols.

**Figure 2.**
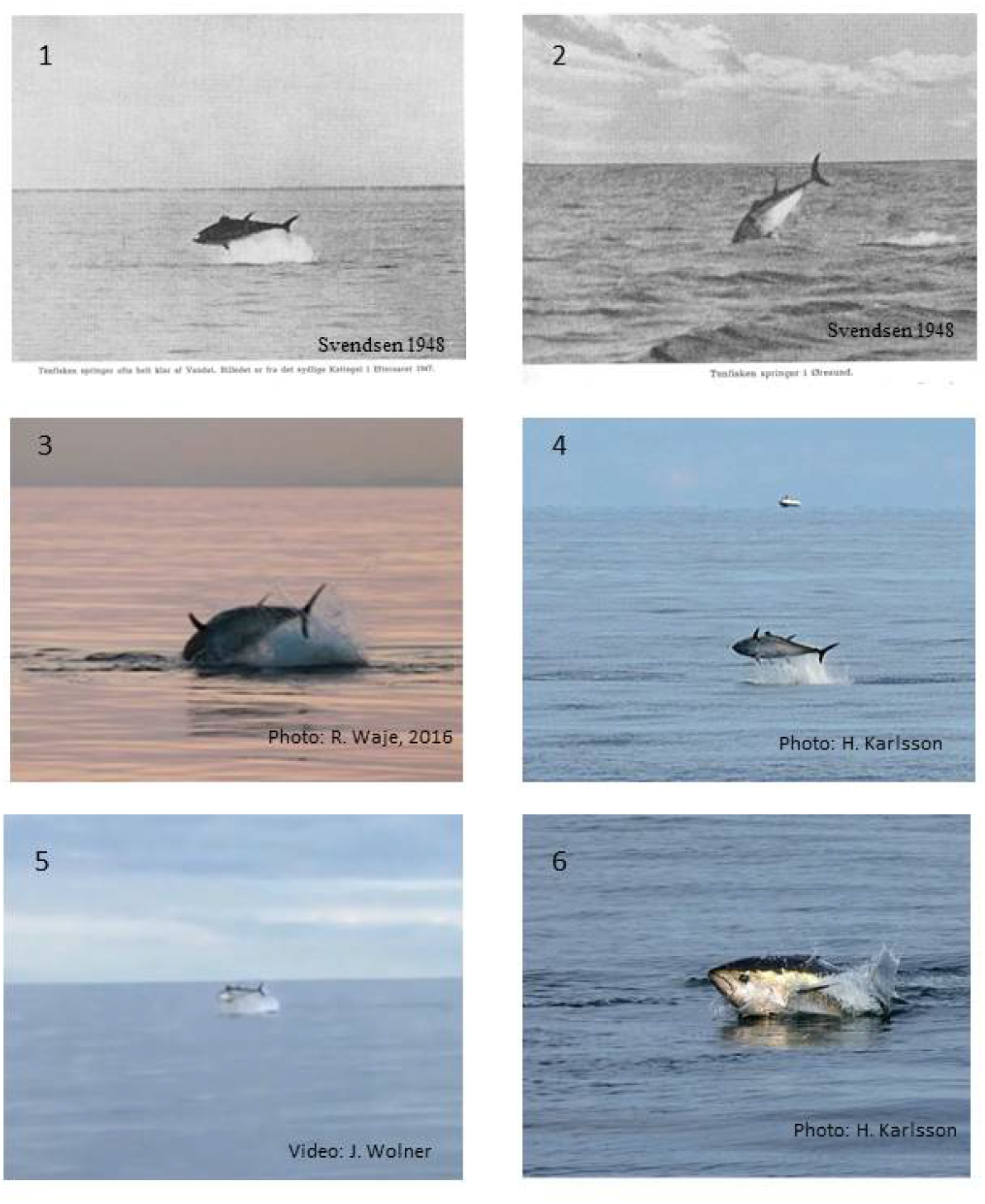
Photographic documentation of presence and jumping behaviour of bluefin tuna in the Skagerrak and Kattegat in a historical period (1947 (33)) and in 2016 (four lower photographs). The historical photographs illustrate the similarity of historical bluefin tuna size, shape and behaviour with that observed in 2015-2017. Additional photographs and links to web-based video clips on social media are available in Supplementary Figure S3 and Table S3. All images are reproduced with permission from photographers.

Our observations were made in 2015-2017. Most observations occurred in 2016 and 2017, although we have records of sightings of 100s-1000s in 2015. The locations of most of the sightings we obtained were in the central part of the eastern Skagerrak, along the Swedish west coast in the Skagerrak, and in the Kattegat all the way down to the Øresund (Figure 1). For example, recreational fishing tour boats, individual recreational fishermen and the Swedish Coast Guard observed individual bluefin tuna swimming at the surface and jumping clear of the water near a Swedish national marine park (Kosterhavet) in 2016. One bluefin tuna was caught by a Danish recreational angler north of Skagen, Denmark in the Skagerrak on Sept. 19, 2016 and released alive after capture. This fish was measured in the water to be 3.03 m and estimated to weigh app. 400-450 kg (25).

At nearly the same time (Sept. 17, 2016) and ca. 7 deg. latitude farther north, bluefin tuna were captured as part of a targeted commercial fishing operation in the Norwegian Sea, off Ona (62.8603°N 6.5543°E), Møre, Norway, approximately halfway between Bergen and Trondheim. Each of the tuna caught (N = 190) weighed ca. 170-300 kg (26–28). Several Danish commercial fishermen, including one of the co-authors of this investigation (HSL), reported that they repeatedly saw schools in a localized area north of Skagen on several days during ca. two weeks in mid-late September, 2016; the total number of tunas observed on some days was in the hundreds. During 2017, we had many more observations than in 2015 or 2016. Several of the observations in 2017 were made as part of a bluefin tuna capture-tag-release project conducted in the area where many bluefin tuna were seen in 2016. During the 8 fishing days, bluefin tuna swimming at the surface or jumping were seen by anglers and/or scientists every day; several anglers, many of whom are professional fishing guides or commercial fishermen for other species, also reported seeing them at depth on their echosounders. Several bluefin tuna were also captured and reported as bycatch by commercial fishing vessels in the Skagerrak-Kattegat in August-September, 2017 (Figure 1).

These 2017 sightings and bycatches in the Skagerrak-Kattegat are supported by other presence data from neighboring areas. These data include commercial catches by Norwegian vessels along the Norwegian coast during August 2017 (29), a bycatch by a Danish fishing vessel 50-60 miles east of the Shetland Islands (North Sea) on Aug. 5-6, 2017 (30), and a stranded bluefin tuna (2.2 m long) found on the Dutch coast on July 31, 2017 (31).

The aggregate seasonality of our presence data in the Skagerrak-Kattegat was from August 23 (2017) – Dec. 4. (2016). The cumulative amount of reports indicates that the species was present in large numbers for at least 2 months during late summer-autumn.

### Ecosystem conditions

The longest available time series of potential prey biomasses are from ICES stock assessments for herring and mackerel stocks in the North Sea, Norwegian Sea and northeast Atlantic. These data show that the biomasses have been high since the late 1980s-early 1990s for these three stocks in the northeast Atlantic Ocean. The sum of the three biomasses has been at record-high levels since the early 2000s (Figure 3).

**Figure 3.**
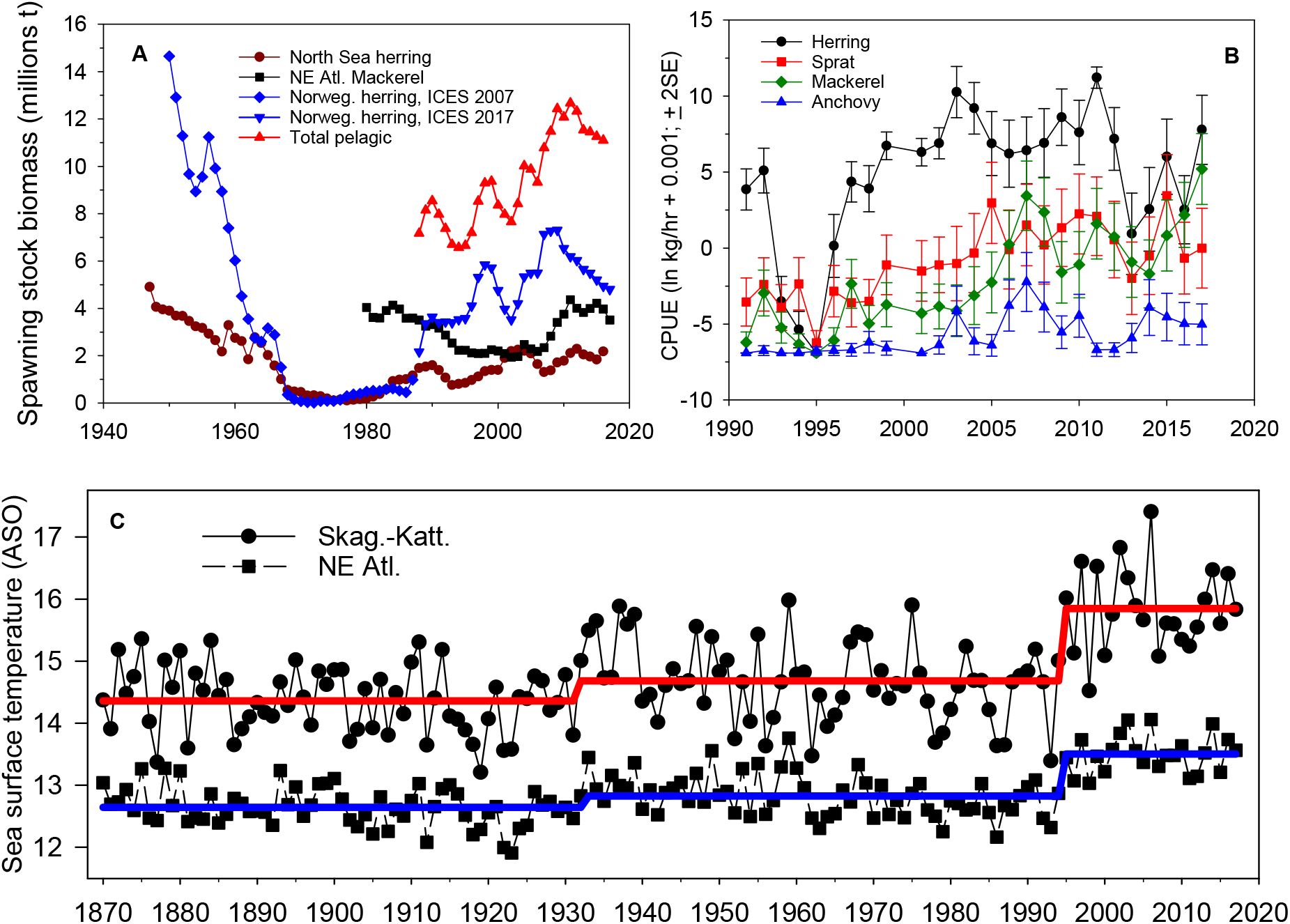
Indicators of potential prey biomass and temperature conditions in the Skagerrak-Kattegat and neighboring regions. A: interannual variability in spawning stock biomass of key prey species (and their sum) for bluefin tuna in northern European continental shelf regions. The stocks are autumn-spawning herring in the North Sea, spring-spawning herring in the Norwegian-Barents Sea, and mackerel in the northeast Atlantic (44). B: mean ln CPUE (ln kg/hour + 0.001; ± 2 x standard error) for herring, sprat, mackerel and anchovy in the Skagerrak-Kattegat during August-September research vessel surveys (International Bottom Trawl Survey, IBTS). C: inter-annual variability in late summer-autumn (mean of August, September and October) sea surface temperature in the Skagerrak-Kattegat and a larger area of the northeast Atlantic (20° W – 13° E; 50° – 66° N) for 1870-2017 (black solid line with dots and black dashed line with squares, respectively) and regime-specific mean temperatures (red: Skagerrak-Kattegat; blue: northeast Atlantic) for different statistically significant regimes (32). Data source: Hadley Climate Research Unit, UK (65)).

At the smaller spatial scale of the Skagerrak-Kattegat, demersal research surveys in late summer-autumn show that catch rates (considered to be a relative indicator of biomass) for four potential prey species (herring, sprat, mackerel, anchovy) were relatively low in 2016 and 2017. The most abundant of these species is usually herring; however its abundance peaked in 2011 and has declined to low levels since then, including in 2015-2017. Other demersal and the pelagic surveys also show that prey abundance in 2015-2017 was approximately average or even below-average (Supplementary Figure S4).

August-October surface temperatures have been well above long-term average since 1994 when a significant regime shift occurred (Figure 3; STARS test; P < 0.0001 (32)); this shift is evident in the larger northeast Atlantic region and the Skagerrak-Kattegat sub-region where our tuna sightings have been made. However temperatures during 2015-2017 were not unusually warm compared to other years during the most recent regime.

## Discussion

Bluefin tuna appear to have re-entered former foraging habitat in northern European waters, which they vacated about 50-60 years ago. The observations are identical to those reported in historical fishery reports, newspapers and scientific literature from the 1920s-1960s (Figure 2), when bluefin tuna were common in these waters (e. g., (33, 34)); the jumping and surface-breaking swimming behaviour is typical for bluefin tuna foraging on prey species (33, 35). Our observations indicate that bluefin tuna were abundant throughout the combined Skagerrak-Kattegat and coastal Norwegian Sea region during especially 2016-2017. Since neither Denmark nor Sweden has a fishing quota, and there are no surveys potentially monitoring the distribution and abundance of bluefin tuna, the public observations are essential for providing evidence of their return to these waters.

Our observations in the Skagerrak-Kattegat during these three years are supported by and consistent with many other reports of bluefin tuna occurring in northern parts of the northeast Atlantic since at least 2012. In that year, bluefin tuna were observed for the first time known to science in waters off east central Greenland (Denmark Strait) as bycatch during mackerel fishing (36). Bluefin tuna were also caught as bycatch in that area in subsequent years (37, 38). Bluefin tuna have also occurred more frequently in waters near Iceland and Norway since the early-mid 2010s. The increased abundance in these waters has led to re-establishment of commercial fishing quotas (2014 onwards) and catches for Iceland and Norway (39). For example, one Norwegian fishing vessel captured 21 bluefin tunas as bycatch in September 2015 in the Norwegian Sea (ca. 80 nautical miles NW of Ålesund) (40). We noted that during the same year and month, large numbers of bluefin tuna were also seen in the Skagerrak-Kattegat by citizen scientists, including one individual bluefin tuna found stranded on the Swedish west coast (Figure 3). In 2013, one dead bluefin tuna (2.2 m long) was found on a Dutch beach (41); prior to this stranding and the one in 2017 (31), no bluefin tuna had been reported stranded on Dutch beaches since 1920 (41) when tunas were more common in Dutch coastal waters (42). The collective volume of reports of bluefin tuna in the wider northern northeast Atlantic (i. e., from Sweden westwards to east Greenland) indicates that the species has been present since at least 2012 onwards, with a documented re-appearance in the Skagerrak-Kattegat starting in 2015. The return to this region, and more generally the northern part of the northeast Atlantic, is therefore not likely a local, short-term anomaly, but more consistent with, and part of a widespread increase in range since at least 2012. Below we discuss potential reasons for this change in spatial distribution.

### Cause of reappearance

Given the long absence from the Skagerrak-Kattegat and neighboring waters, one must ask why bluefin tuna has finally returned now. We believe that the factor which has likely contributed most to the return is improved bluefin tuna fishery management since ca. the mid-late 2000s. Several changes were made during this period, including reductions in quotas, increases in minimum landing sizes from 6 to 10 kg and then to 30 kg so that a much larger share of juveniles can now survive long enough to reach maturity (assumed to be at an age of 4 years, or ca. 25 kg (12)), improved catch reporting requirements and documentation, a shorter fishing season during the main spawning period in the Mediterranean Sea and strengthened fishery surveillance and enforcement (12, 13).

Prior to implementation of these changes, the stock was overexploited both legally (because ICCAT’s contracting parties were adopting total allowable catches higher than those recommended by ICCAT’s scientific body [Scientific Committee on Research and Statistics, SCRS] as being scientifically sustainable in the long term), and illegally (e.g., landings often exceeded, sometimes even by 2-fold, the quotas adopted by the contracting parties during many years in the late 1990s and early 2000s). Before the new regulations were implemented, historical exploitation of juveniles in southern parts of the stock range was high since the 1950s and has been considered to be a major factor leading to the disappearance and subsequent continued absence of bluefin tuna from northern parts of the range (20). In addition, the parent stock biomass in the early-mid-2000s was perceived to be declining (12) and at a rate that if continued could have met criteria for listing this stock as “critically endangered” according to IUCN (International Union for the Conservation of Nature) criteria (43).

Implementation of the new fishery management regulations, and a real and effective monitoring control and surveillance system, appears to have had positive effects on the Mediterranean stock of bluefin tuna. Shortly after, several stock indicators of abundance started increasing, including the biomass of adults and the production rate of new young bluefin tuna “recruits” (12, 13). Moreover standardized catch rates of adults and juveniles in several regions of the northeast Atlantic and Mediterranean Sea by commercial fishing vessels have increased since the implementation of these new regulations (39).

As the stock has increased, bluefin tuna appears to have expanded its migratory range, a pattern common among recovering fish stocks (3, 22), to explore new feeding habitats and to reduce density-dependent competition for prey, including into some northern areas beyond formerly documented distribution ranges such as Denmark Strait (east of Greenland (36)). This exploratory foraging behaviour may have led them to return to the northern European shelf waters, including the Norwegian Sea, North Sea and Skagerrak-Kattegat where they apparently have found sufficient prey for foraging.

A second reason for the return to these waters may be the relatively high biomasses of potential energy-rich prey species such as mackerel, herring, and sprat. Herring and mackerel dominate pelagic fish biomass in the northern part of the northeast Atlantic Ocean and their biomasses have recovered to or beyond historical estimates (Figure 3)(44). Some of these species (herring and mackerel) were also overexploited in the 1950s-1970s leading to local collapses and fishery closures for these species. These declines may have been a factor inhibiting earlier return of bluefin tuna to this region. However following implementation of more sustainable fishing practices, the biomasses of these species recovered to long-term average levels by the early-mid 1990s and have remained high or increased further since then (Figure 3). However, the tunas did not reappear in large numbers until the early-mid 2010s (i. e., ca. 10-15 years after the prey biomass indicators recovered). This delay coincides with the period when bluefin tuna were exploited at levels well beyond those considered sustainable and when overall biomass was declining, leading to development and implementation of a recovery plan (12, 45, 46). The delayed return, despite high prey biomass in the North and Norwegian Seas, suggests that the main reason for the reappearance of bluefin tuna in the region was the increase in tuna biomass itself, and the time required for individuals to re-learn former migration pathways and foraging habitats (19, 47). It is possible therefore that if bluefin tuna had been more abundant in the 1990s and 2000s, they would have already appeared in high abundance in the Norwegian-North Sea-Skagerrak-Kattegat area several years earlier than now.

Ocean temperature conditions are also known to affect tuna distributions and migrations (18, 48, 49), and have been generally warmer since 1994. However temperatures in 2015-2017 were not exceptionally warm and were in fact colder than in some earlier years in the recent warm regime (post-1994). Notably, bluefin tuna have been present in the Skagerrak-Kattegat during many earlier years (e. g., 1920s-60s) when temperatures were lower than during the post-1994 regime (compare Figure 3 and Supplementary Figure S2), and have occupied colder areas farther north in the past (e. g., in the Norwegian Sea and along the west Norwegian coast; see Ref.(34) for temperature data). We conclude that temperatures in the Skagerrak-Kattegat or northeast Atlantic region appear to have had little direct role on the re-appearance of bluefin tuna, although they may have had indirect effects via changes in abundance, migratory behaviour or distribution of local prey species. However the exact mechanisms by which temperature may have acted are unclear and remain speculative.

In summary, both food and temperature conditions have been higher than average for many years before the tuna returned to the Skagerrak-Kattegat and probably also the wider North Sea-Norwegian Sea-Skagerrak-Kattegat region. We consider it unlikely that either of these factors in 2015-2017 were the main direct drivers for the recent appearance of bluefin tuna in the Skagerrak-Kattegat region, although adequate prey and temperature conditions likely induced exploratory foraging bluefin tuna to remain in and potentially return to this region, once it was re-discovered. Consequently local stocks of prey species such as the western Baltic spring-spawning herring, which migrate through the Skagerrak-Kattegat when bluefin tuna are seasonally present, probably stimulate bluefin tuna to remain and forage in the region after they have arrived and explored in the Norwegian-North Sea region.

The recent increase in overall bluefin tuna biomass in the northeast Atlantic-Mediterranean Sea has probably led to an increase in exploratory and more distant foraging behaviour by some individuals. Some of these individuals and schools of bluefin tuna have apparently encountered the larger biomass of potential prey species, including mackerel and herring in the region, and have in turn transferred or shared migratory and habitat knowledge with other individuals. These mechanisms are known to be key elements of the discovery of new habitats and connectivity among geographic regions in migratory populations (5, 19, 21, 23) and may have led to the reappearance of bluefin tuna in the Skagerrak-Kattegat and more generally in the northern northeast Atlantic, including towards east Greenland and Iceland (36), where mackerel biomasses have been increasing (50, 51). New tagging studies in the region will help address these mechanisms in future.

Although the mechanism by which bluefin tuna have learned to migrate to the Skagerrak-Kattegat-North Sea-Norwegian Sea region is unclear,these newly-acquired feeding migrations may have direct benefits for the population spawner biomass, egg production and overall sustainability of the stock. Historical feeding in the North and Norwegian Seas allowed bluefin tuna to rapidly increase their overall condition factors due mainly to consumption of energy-rich prey species (17, 52). Fish in better condition typically have higher and better quality reproductive output, potentially leading to higher recruitment per spawner and are less likely to skip spawning (53). Development of new annual feeding migrations to this area may therefore have benefits for population fecundity and recruitment.

### Future perspectives

The return of bluefin tuna to the Skagerrak-Kattegat in 2015-2017 seems to be part of a longer-lasting seasonal re-occupation of these and other northern European waters. If this situation continues in the future, it will open many new possibilities for both scientific understanding of species biology and dynamics, and for socio-economy opportunities as for example sustainable fisheries (commercial and recreational), catch and release angling, and eco-tourism. Regarding science, a priority should be to investigate the migration behaviour and population origin of the bluefin tunas which have appeared in these waters using a combination of modern tagging, genetics, otolith and modelling methods. Bluefin tuna in the Atlantic are managed as two stocks (western and eastern stocks, the latter including the Mediterranean) with separate quotas and regulations (12). Historically, bluefin tuna caught in the northern European region had migrated both from the northeast and northwest Atlantic (14), although the historical proportions are not known with certainty. Any future commercial or recreational catches should be assigned to the correct stock. Such assignment would need for example genetic or otolith-based evidence (54, 55). New sustainable commercial and recreational fisheries would support and diversify local fishery economies in primarily rural areas of the region. Furthermore, given the recent interest in the general public for the return of this species to northern European waters (56, 57) (e. g., 7 video clips made by citizen scientists have been viewed on social media > 510,000 times since September 2015: Supplementary Table S3), and the possibility that jumping bluefin tuna can be seen from relatively small boats operating within minutes to a few hours of shore, there is potential that the species could create and contribute to the eco-tourism industry.

A pre-requisite for realizing these scientific and socio-economic opportunities is that the recent fishery management regulations for both bluefin tuna and their prey, and their compliance, continues in future. In general, it presently appears that efforts in the past decade towards sustainable fishery management both for the bluefin tuna and its prey species are having positive benefits for these stocks. Should this be true, bluefin tuna may become a regular summer component of local fish communities and food webs (as it historically has been; (52)), and contribute to small but lucrative commercial and recreational fisheries and eco-tourism economies. If our working hypothesis that the reappearance is due mainly to recovered bluefin tuna and prey stocks, then management actions which ensure a modest-high biomass of large individuals in the tuna population will be needed.

We note that commercial fishing quotas for the NE Atlantic-Mediterranean stock will increase gradually until 2020 by 13,295 t or 58 % relative to the 2017 quota (58). The population abundance estimate used as basis for this increase has many uncertainties of unknown magnitude (39, 59, 60). As a result, should the stock not be, or grow, as large as expected, then the higher quotas could lead to a population decline. If this happens, then it is possible that the abundance of the larger individuals containing the migratory memory and knowledge will decline; this could lead to a retraction from the northern area again. We further note that the size range of bluefin tunas seen in the area in 2015-2017 is relatively narrow (generally 2-3 m), suggesting that the migrants which have arrived during 2015-2017 are from a relatively small number of year-classes born in the early-mid 2000s (assuming a stock-specific length-at-age relationship (61)). If the number of year classes observed in northern European waters is indeed small, and if post mid-2000 year classes are weak-average, then the combination of ongoing natural mortality, higher landings and fewer new recruits could lead to reduced immigration to northern European waters in the future. Continued monitoring of stock size, distribution and habitat characteristics (e. g, prey abundance and size), including tagging, is needed to improve our understanding of factors affecting bluefin tuna ecology in the region.

The re-establishment of summer migration to the Skagerrak-Kattegat, together with the recent increase in overall stock biomass, indicates that, as with some other recovering large iconic fish stocks (3), improved management, enforcement and compliance can yield positive benefits when ecosystem conditions for stock production are suitable (3, 9). These observations apply even for species such as bluefin tuna which has historically suffered from much illegal and over-fishing and whose highly migratory behaviour takes them into multiple fishing jurisdictions including international waters. The reappearance of bluefin tuna in the Skagerrak-Kattegat and neighboring seas (e. g., North Sea, Norwegian Sea) serves as an example of the benefits of implementing effective recovery actions, despite a decades-long absence from the region and a highly unsustainable fisheries exploitation situation. In this case, implementation has not been too late to promote recovery. Similar efforts with other populations and species could also yield positive outcomes. These findings, as well as some other recent and ongoing major marine recoveries (3, 62, 63), offer some optimism for the long-term recovery and sustainability of commercially-exploited fish stocks, the ecosystems in which they live, and the economic sectors which they (could) support.

## Materials and Methods

### General

As there are presently extremely limited, if any, commercial or recreational fisheries (i. e., Denmark and Sweden have no quota, but Norway does) for bluefin tuna in the Skagerrak-Kattegat and no scientific fisheries (surveys) for this species in the region. Therefore the main source of information available for documenting presence was reports from the public, including commercial and recreational fisher observations. We consider all these records to be “citizen science” and compiled observations from reports in newspapers, social media and via direct contact of the public with us. Details for the data compilation are available in Supplementary Material. We supplemented the citizen science observations with our own observations in 2017 as part of a dedicated tagging project in the area (24) (observational details in Supplementary Material). All observations were then organized chronologically and visualized to display their spatial-temporal distribution. To place our work in the historical context of past fisheries for bluefin tuna in the region, we compiled and plotted officially reported landings data from ICES and other historical sources (summarized in (34)) for the years 1903-2014, which was the last year for which landings data are available from ICES. We use officially reported data from ICES instead of from ICCAT because the former are available at higher spatial resolution for our region of interest.

Ecosystem conditions (e. g., local food concentrations, temperature) are known to affect distributions of bluefin tuna (18, 36). We derived estimates of the inter-annual variability in abundance of major prey species and sea surface temperatures (i. e. those most likely experienced by bluefin tuna during summer foraging on the continental shelf) for the northeast Atlantic to evaluate whether food and/or temperature conditions were unusually favorable in 2015-2017 compared to earlier years. Further details of the data sources and compilation are presented in Supplementary Material.

## Acknowledgments

We thank Jeppe Olesen for GIS and mapping expertise and members of the public without whom this investigation could not have been possible. The work was partly supported by the Nordic Council of Ministers (project no. 141-2016-Tuna; NordTun), ICCAT (Tagging programme on bluefin tuna GBYP Phase 7) and has been conducted using MyOcean Products.

## Supplementary Materials

### Supplementary Text

#### Methods

This section provides details how bluefin tuna observations were compiled, how they should be interpreted, and the sources and processing of ecosystem data (prey abundances, sea surface temperatures)

#### Bluefin tuna observations

We compiled observations of bluefin tuna from primarily public sources such as social media (i. e., Facebook, YouTube) and websites representing both commercial fishermen and anglers in Denmark and Sweden. We also obtained and used information sent to us by the public following announcements on the DTU Aqua website in 2016 and 2017 and sent to Danish commercial and sportsfishermens’ organizations and contact by SLU Aqua with Swedish sportsfishermen that we were interested in receiving sighting observations. We supplemented these citizen science observations with reports of commercial catches and bycatches in the Norwegian Sea, North Sea, Skagerrak, Kattegat and Øresund. The time period covered was August-October in 2016 and 2017. However during the course of this data collection, we received or found reports of observations in 2015 which we used to support our overall results and conclusions. In many cases, the sightings were supported by photographs or video recordings of bluefin tuna. The information provided by members of the public included date and location of the observation, how many bluefin tuna were observed (e. g., single individual, school, approximate number of schools and number of fish per school), behaviour (jumping over water surface, breaking water surface), and prey escape behaviours observed at the surface. The observations were entered into a database and visualized geographically to illustrate their spatial distribution in relation to distance from land, bottom topography and sea surface temperature. We are aware that the citizen science data may not necessarily represent the true spatial distribution of the bluefin tunas; details of the data collection and their interpretation are presented in Supplementary Material.

We also made our own observations in 2017. These observations were gathered during a bluefin tuna tagging study conducted in the eastern Skagerrak-Kattegat. Fishing operations were conducted on 8 days in September, 2017 by ca. 30-40 angling boats. As part of the tagging study, we requested anglers to record sightings of bluefin tuna jumping and swimming at the surface. These sightings, along with the hooked, tagged and released, and hooked-escaped bluefin tunas were used in our compilation. Full details of the tagging operation are described elsewhere (1).

#### Contribution of citizen science to bluefin tuna ecology

For reasons explained in the text, our investigation relies heavily on input from the public to document the species presence in this region. As with all citizen science reports, there is a possibility for some false, biased and otherwise incorrect reporting. We believe that such records are not likely present in our compilation because of the nearly simultaneous nature of the records over a wide area (e. g., the many sightings reported in the eastern Skagerrak-Kattegat on nearly the same day in 2016 as a large commercial catch in central Norwegian coastal waters (2); the sightings, capture, tagging and release of bluefin tunas in the eastern Skagerrak-Kattegat during September 8-24, 2017 (1) and a commercial catch of 78 tunas in Norway in late August, 2017 (3)), the similarity of the reported behaviour to historical sightings of tuna in the region (e. g.,(4); see Figure 2 of main manuscript and Supplementary Figure S3) and our own observations during the 2017 tagging operation, and the distinguishable features of bluefin tuna behaviour and size that reduce the likelihood of misidentification with other species.

Moreover, some of the citizen science reports were made by highly reputable observers, including on-duty officers of national coast guard services or off-duty members of our research vessels while participating in recreational sea-based activities. Their observations and reports were identical to those made by other members of the public, by commercial and recreational fishermen targeting other species and by ourselves during our 2017 tagging study. For example, some fishermen observed tuna while trawling for pelagic fish (e. g., herring and mackerel) that are prey for bluefin tuna in this region. Lastly, several of the reports and our interviews via email or telephone include statements by the observers that they had never seen such behaviour before despite years and even decades of activity on the sea and that they had knowledge of the species’ former presence in the area from older generations. We are confident therefore that our observations represent a valid and reliable source of documentary evidence of the presence of the species in these waters.

We are aware however that the reports based on citizen reporting reflect the spatial distribution of where the reporting observers were located, and also our own efforts to obtain such information, which started in 2016. As a result, such data do not necessarily represent the full spatial distribution of where the tuna were located because (1) some tuna may have been observed in other areas, but not reported to us, (2) some tuna may have been present in other areas (and of course other depths), but not seen by any human observers; and (3) observers were surely present over a much wider area than indicated by our few reports, but it is not possible to know which of those observers saw or did not see bluefin tuna. The spatial and temporal distribution of tuna based on our observations must therefore be interpreted cautiously, and we cannot exclude the possibility that bluefin tuna were present over a much wider area or at other times than is indicated by our data. We have tried to minimize such observer bias by making broad contact to the public and especially commercial fishermen (e. g., via their associations). Nevetheless, to obtain a more representative distribution in the area, alternative methods would need to be employed such as acoustic surveys, aerial surveying via airplanes (5, 6) or with drones (7) or tagging with electronic tags (8, 9) and subsequent modelling (10, 11). In addition, increased public awareness of the species in the area and the need for its documentation could also increase public reporting of bluefin tuna observations and the reliability of distributional maps. Such combined survey-citizen science methods could also potentially be used to derive estimates of relative abundance in the region, which is not possible with our dataset.

#### Estimates of abundance of potential prey for bluefin tuna

We estimated abundances of potential prey for bluefin tuna in the region from regional stock assessments for main prey species and from fishery research vessel surveys. We used the North Sea herring, Norwegian spring-spawning herring and Northeast Atlantic mackerel total stock biomass estimates from the ICES assessments as indicators of potential prey for bluefin tuna in the region of our study. These stocks occupy large areas (see ICES stock management area map, Supplementary Figure S1), which overlap with the historical distribution of bluefin tuna in the region (12–15); moreover tuna which enter the Skagerrak and Kattegat historically passed through the northern North Sea and Norwegian Sea on their way to this region (12, 14) and would potentially encounter these prey during the migration and while foraging for prey. These biomass estimates are based on stock assessments of the various stocks (16).

We also used scientific research vessel surveys to estimate prey abundances more locally in the Skagerrak-Kattegat and where bluefin tuna were observed in 2015-2017. Hydro-acoustic and bottom trawl surveys are conducted annually in the region as part of population status monitoring in the region for fisheries management purposes (17–20). The surveys are conducted in February-March (demersal survey in Kattegat-Belt Sea), late June-early July (hydro-acoustic pelagic survey in Skagerrak-Kattegat), and August-September (demersal survey in Skagerrak-Kattegat). The three surveys when considered in aggregate provide information about the relative abundance of potential prey (herring, sprat, mackerel) among years. The demersal survey in August-September is conducted when bluefin tuna were present in our area and we present its results in the main article; survey estimates at other times of year are presented as Supplementary Information.

The main characteristics of the surveys (depth sampled, geographic location, year and seasonal coverage) are summarized in Supplementary Table S1. The notable feature for all the surveys is that sampling methods and gear within each survey are the same throughout the time periods shown here (17–20). As seen in the Table, only the acoustic survey is directly designed to estimate abundance and biomass of pelagic fish species (e. g., herring, sprat); this survey is used as input to ICES stock assessments for herring in the western Baltic Sea and in the North Sea (17). The other two surveys are designed to capture demersal fish species as part of the International Bottom Trawl Survey (IBTS) (21) and Baltic International Trawl Survey (BITS) (18). However these surveys regularly capture pelagic fish species, including herring and sprat, and these data can indicate relative trends and fluctuations in biomass, even though they may not necessarily represent true abundances due to lower catchability for pelagic fishes which are distributed higher in the water column than the demersal sampling gear. For the acoustic survey, we use the total abundances of all size groups estimated on the survey in specific strata of the survey (i. e., Kattegat, and waters along the Swedish west coast in the northeastern part of the Skagerrak (19)). For the demersal surveys, we calculated annual geometric means of catch-per-unit-effort (CPUE) using biomass/trawl hour as a relative biomass metric. Full details of the survey methods and sampling gear are available in literature (17–20, 22).

#### Estimation of sea surface temperature

Bluefin tuna occupy mainly surface waters (i. e., above the seasonal thermocline) when feeding on continental shelves in summer as in the region of our study. This habitat is also the depth layer predominantly occupied by their main prey (e. g., herring, sprat, and mackerel) in the region. We assumed that sea surface temperature (SST) as estimated by satellite imagery is an approximate indicator of the temperatures available for and experienced by bluefin tuna while foraging in the region.

We calculated the average SST for the region for the months of August, September and October for the time period 1870-2017 using a large international database of in situ and satellite-derived observations(23) available online (http://wps-web1.ceda.ac.uk/ui/home) at 1 degree monthly resolution. This time series allowed us to evaluate whether 2015-2017 were exceptionally warm summers relative to historical variability and trends. We calculated mean temperatures for both the Skagerrak-Kattegat (8° – 13° E; 55° – 59° N) and a larger area of the northeast Atlantic Ocean (20° W – 13° E; 50° – 66° N) through which bluefin tuna migrate when entering the Norwegian-North Sea-Skagerrak-Kattegat. We evaluated whether regime shifts in temperature occurred using the STARS algorithm (24) for a significance level P = 0.05; settings used for testing were Huber parameter = 1 and series length = 10. Higher resolution (0.05 degree daily) satellite-based estimates of SST (25) in the Skagerrak and Kattegat region were averaged temporally over the main period where tuna were observed (7th – 21st September during 2015-2017) and used to characterize the thermal environment in which the fish were observed.

## Supplementary Figures

**Figure S1.**
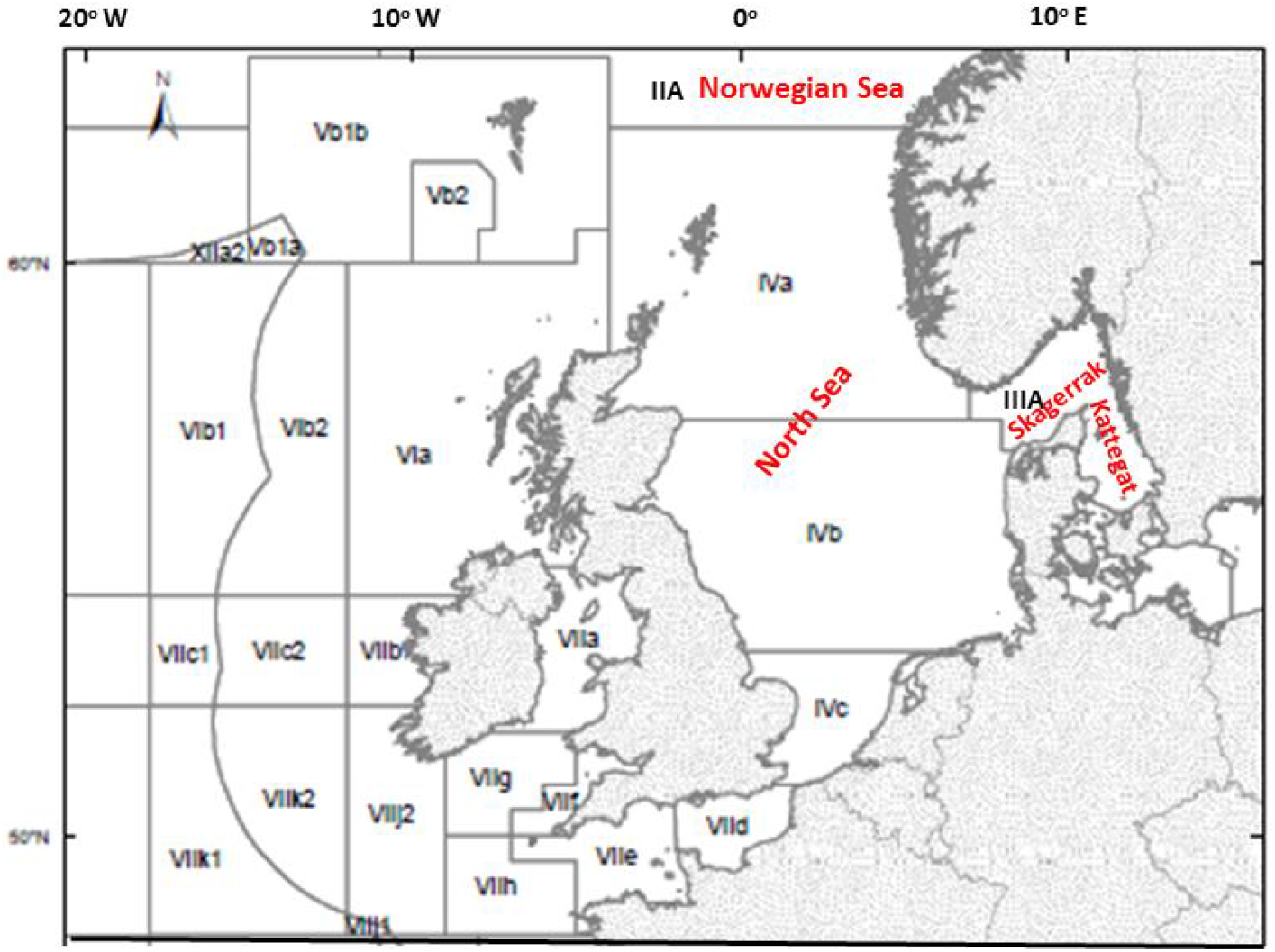
ICES fishery stock management areas.

**Figure S2.**
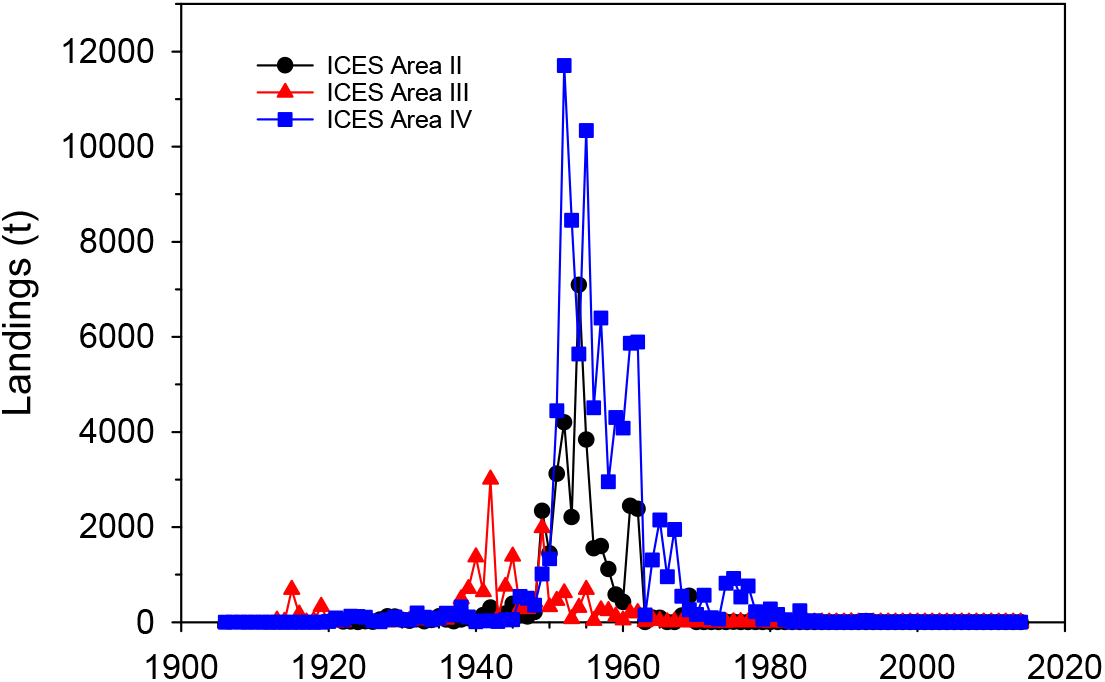
Reported catches of bluefin tuna in ICES areas II, III and IV corresponding approximately to respectively the Norwegian Sea, North Sea, and Skagerrak-Kattegat-Øresund (see Figure S1 for map of ICES stock management areas). Catch data officially reported to ICES from 1903-2014 are from ICES databases available online (www.ices.dk). Additional catch data from before 1927 were compiled from historical fishery reports, catch records, museum records and other documents as summarized in (15).

**Figure S3.**
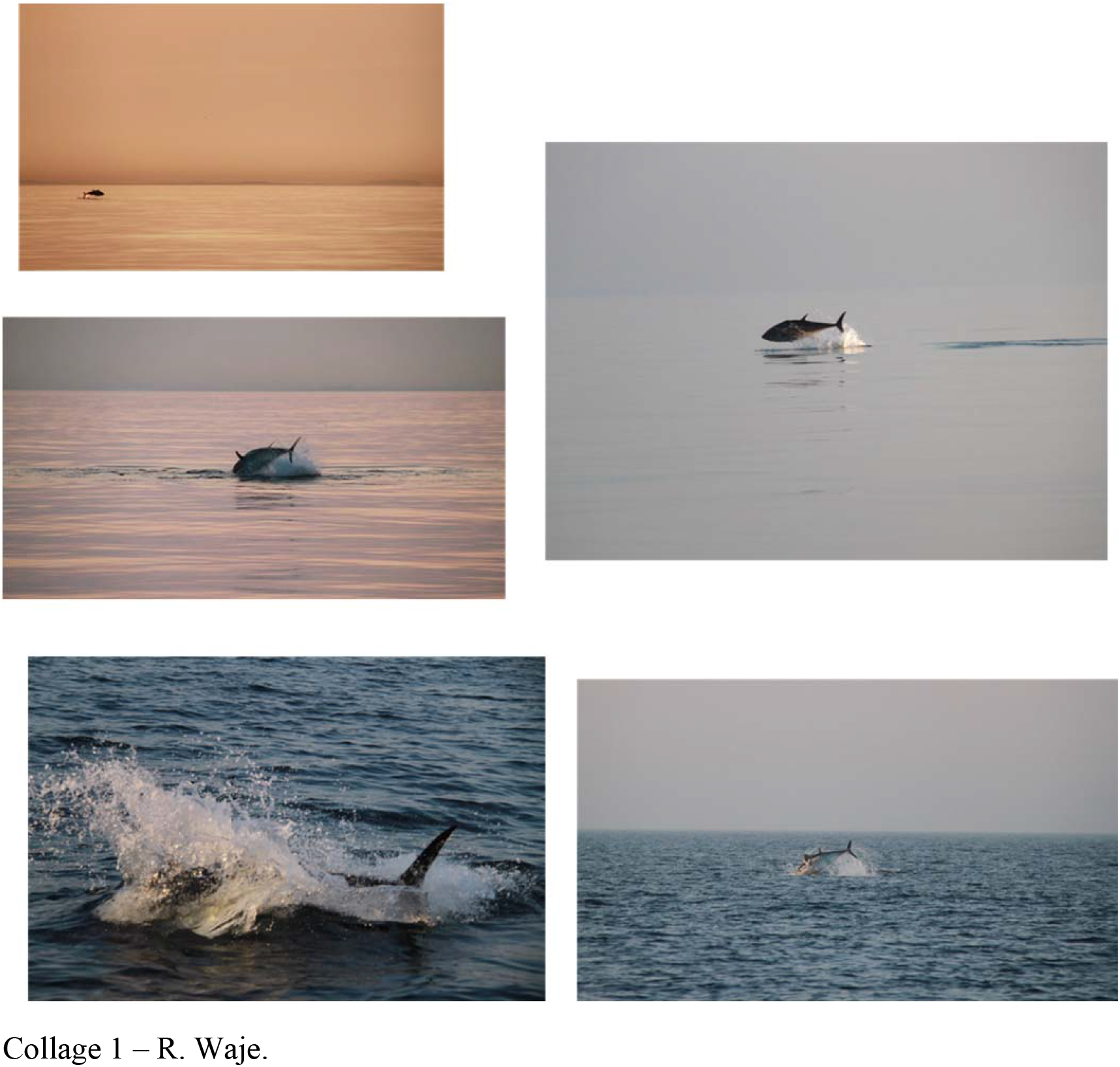
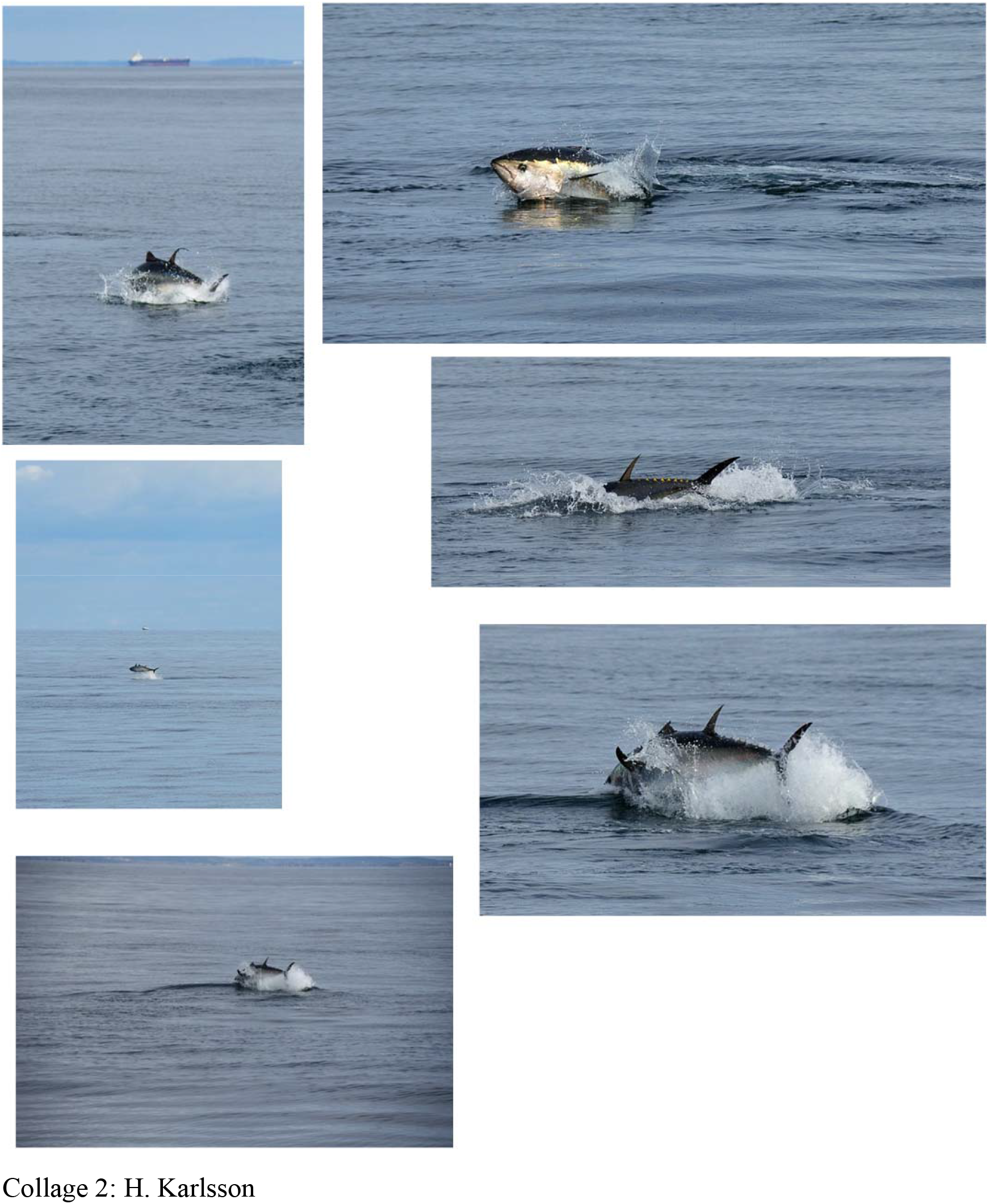
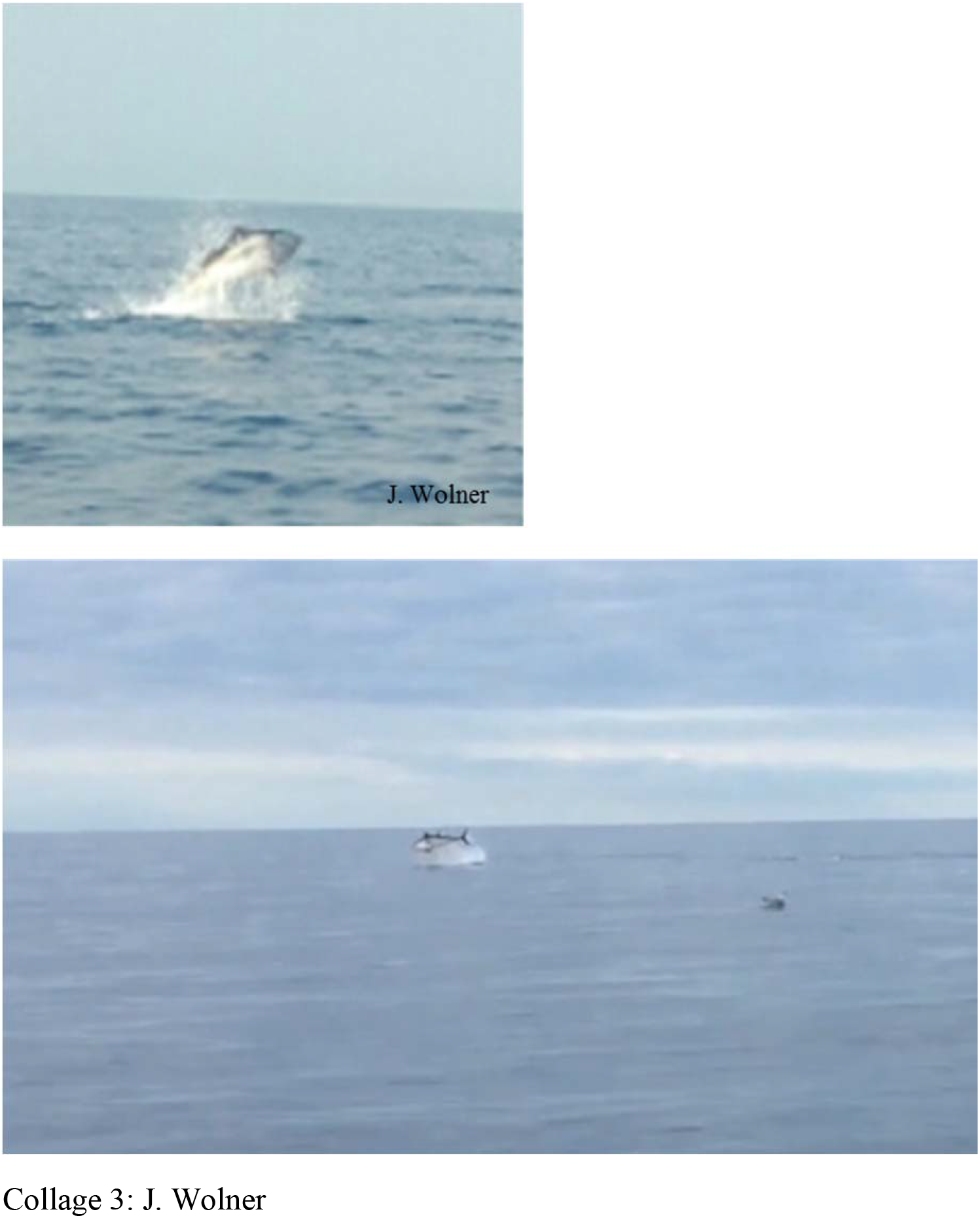
Photographs showing bluefin tuna in the Skagerrak-Kattegat during September 2016. Photos provided with permission by members of the public. Collage 1: R. Waje, collage 2: H. Karlsson, collage 3: J. Wolner.

**Figure S4.**
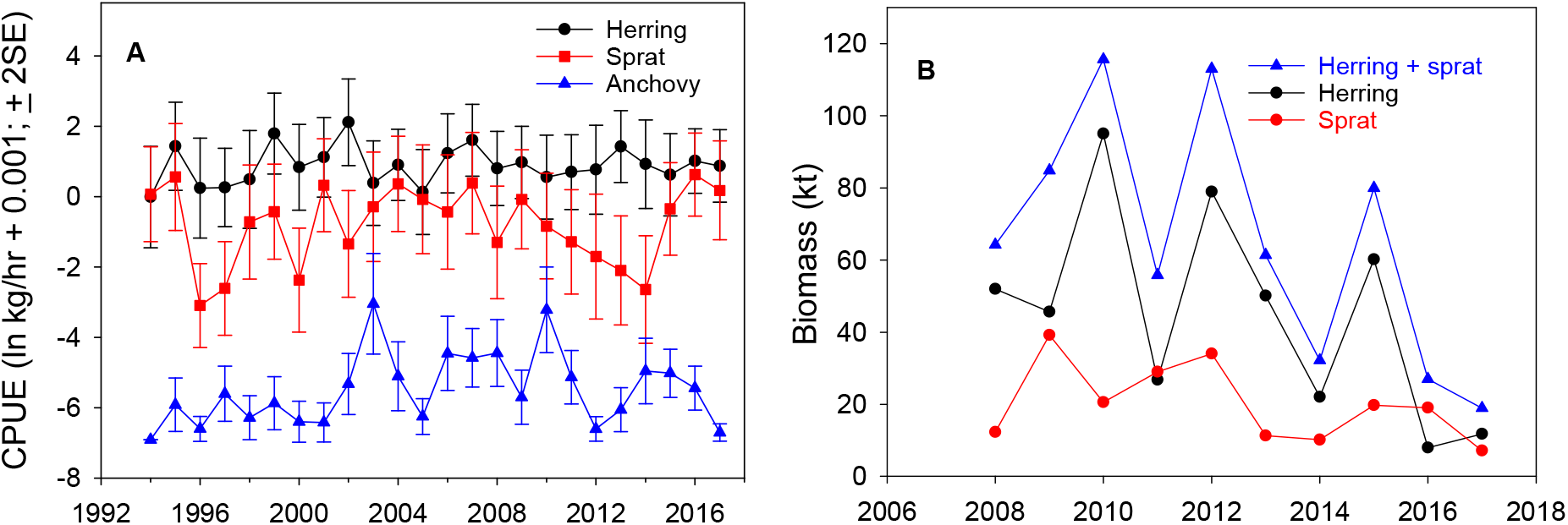
A: mean ln CPUE (ln kg/hour + 0.001; ± 2 x standard error) for herring, sprat and anchovy in autumn (late October-early November) research trawl surveys in Kattegat-Øresund-Belt Sea (north of 55 N.) during 1994-2017. B: estimates of prey biomass as derived from research vessel hydro-acoustic surveys in the Kattegatduring 2008-2017 (see Table S1 for survey details).

## Supplementary Tables

**Table S1.**
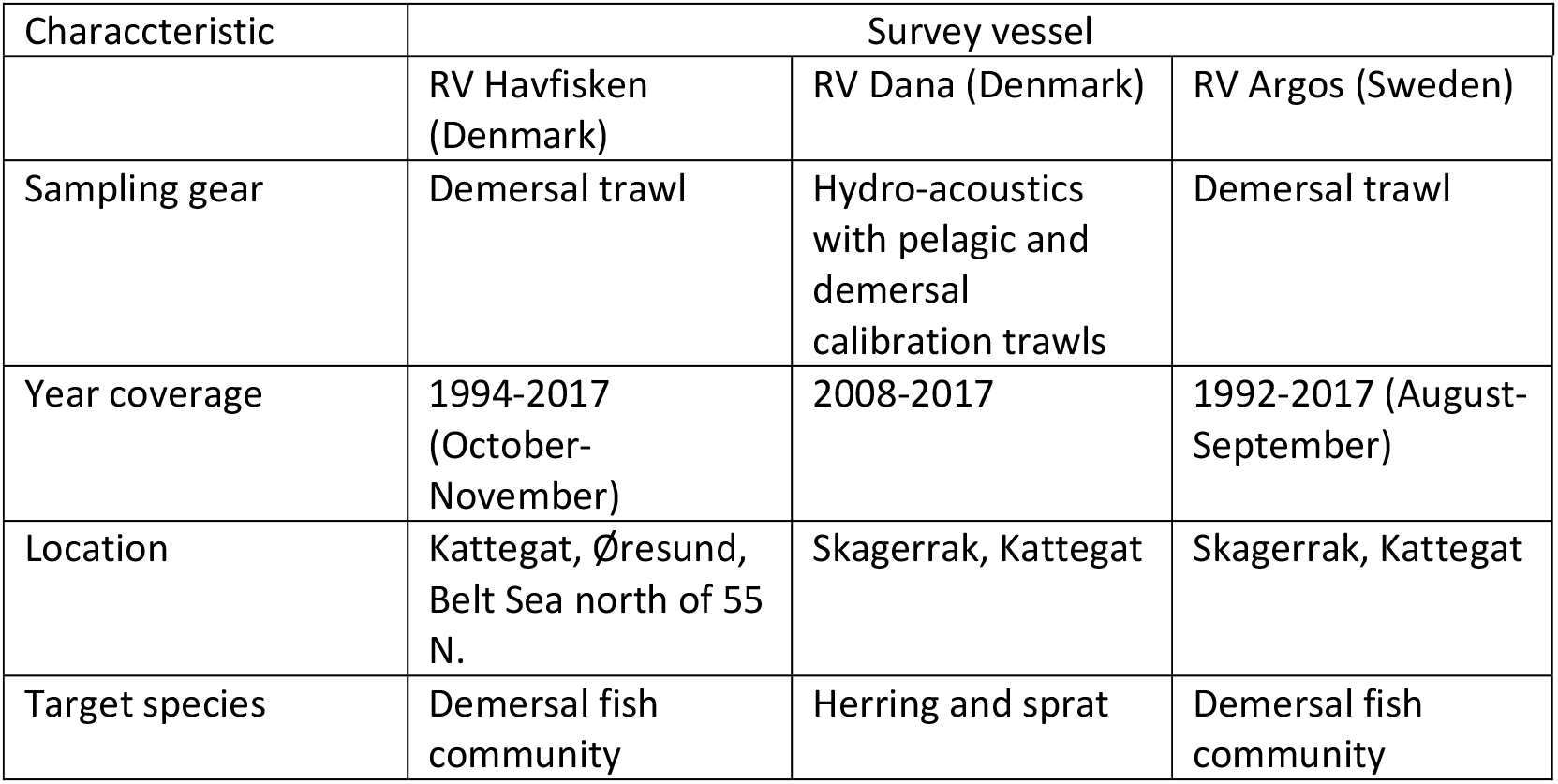
Overview of research vessel surveys which provided estimates of biomass of potential prey for bluefin tuna in the Skagerrak-Kattegat-Øresund. The surveys are part of international bottom-trawl and acoustic surveys in the region designed and coordinated internationally by ICES.

Table S2. Observational data for sightings, commercial bycatches, and research tag-releases of bluefin tuna individuals and schools in the Skagerrak, Kattegat and Øresund during 2015-2017. The data were recorded by members of the public including anglers (“citizen scientists”), commercial fishermen and research scientists. Observers’ identities are known to the authors. Table will be submitted to journal as an excel file supplement.

**Table S3.**
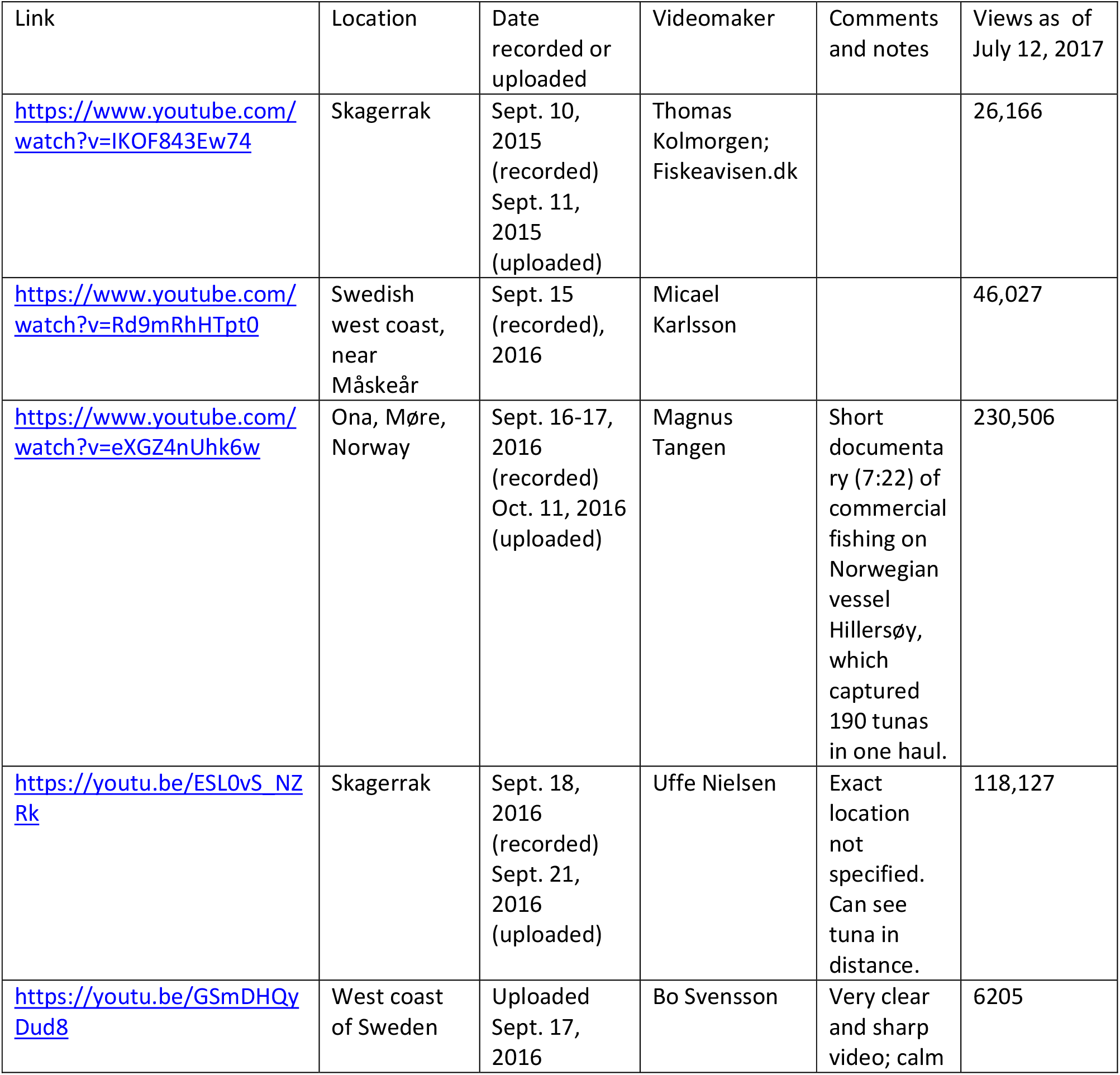
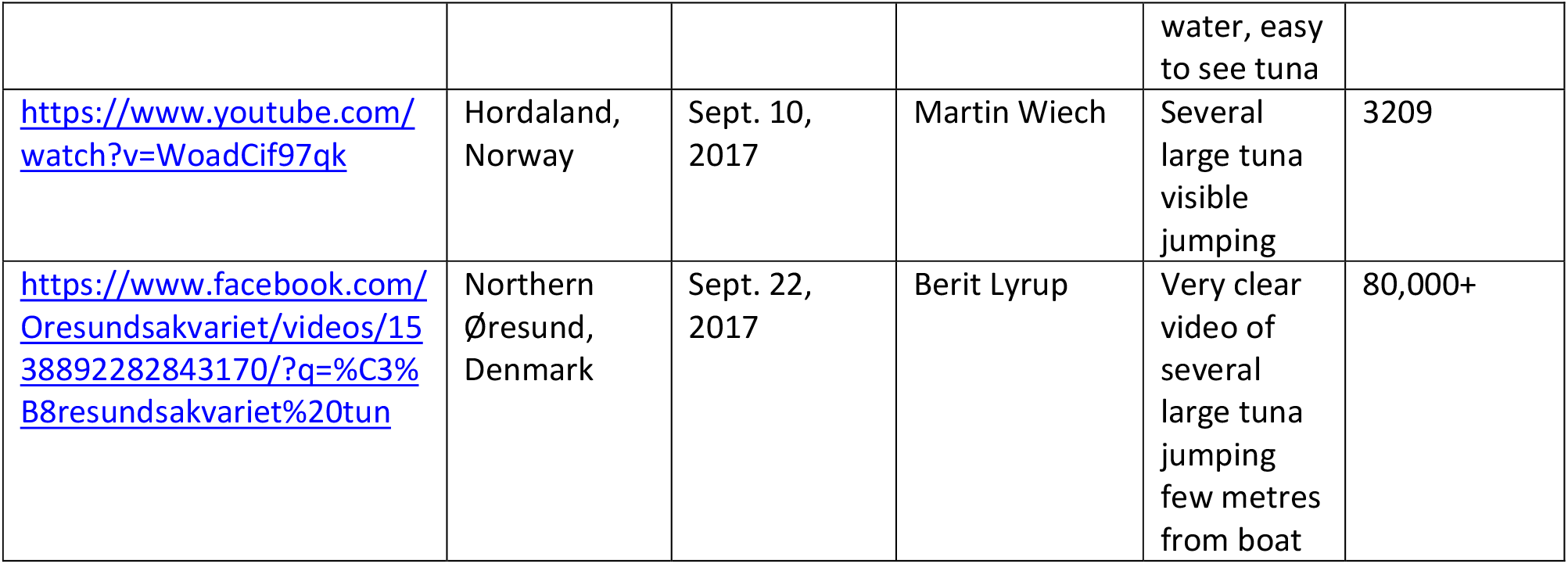
Examples of videos on social media of bluefin tuna *Thunnus thynnus* swimming and jumping at surface in the Skagerrak-Kattegat and off Norway during 2015-2017. Also indicated is the number of views of each videoclip. The total number of views was 510,240 as of April 23, 2018.

## Author Contributions

BRM designed research, compiled data and wrote the manuscript; all authors contributed to the design of the study and/or edited drafts of the manuscript. KA, KBG, MCh, and GQB assisted with data collection in Denmark and MCa, IO and AS assisted with data collection in Sweden. CS and HSL assisted with data collection from recreational and commercial fishers. MRP produced satellite imagery temperature products.

